# MycoPhylo experiment reveals how mycorrhiza types and phylogenetic relationships affect soil biodiversity and functioning

**DOI:** 10.1101/2022.04.26.489578

**Authors:** Leho Tedersoo, Kaire Loit, Ahto Agan, Saleh Rahimlou, Annaliisa Vask, Rein Drenkhan

## Abstract

- Natural forests and abandoned agricultural lands are increasingly replaced by monospecific forest plantations that have poor capacity to support biodiversity and ecosystem services. Natural forests harbour plants belonging to different mycorrhiza types that differ in their microbiome and carbon and nutrient cycling properties.
- Here we describe the MycoPhylo field experiment that encompasses 116 woody plant species from three mycorrhiza types and 237 plots, with plant diversity and mycorrhiza type diversity ranging from one to four and one to three per plot, respectively.
- The MycoPhylo experiment enables us to test hypotheses about the plant species, species diversity, mycorrhiza type, and mycorrhiza type diversity effects and their phylogenetic context on soil microbial diversity and functioning and soil processes.
- Alongside with other experiments in the TreeDivNet consortium, MycoPhylo will contribute to our understanding of the tree diversity effects on soil biodiversity and ecosystem functioning across biomes, especially from the mycorrhiza type and phylogenetic conservatism perspectives.

## Introduction

Forests are the most widespread land cover type globally (Pan *et al*., 2012; Crowther *et al*., 2015) and they harbour much of the biodiversity and soil carbon (Parrotta *et al*., 2012; Scharlemann *et al*., 2014). Forest conversion to agricultural land is one of the most important threats to global biodiversity and a major source of greenhouse gas emissions (Parrotta *et al*., 2012; Wei *et al*., 2014). Plantation forestry has only partly mitigated these concerns from the carbon sequestration perspective but not from the biodiversity perspective (Brockerhoff *et al*., 2008; Nave *et al*., 2013). Monoculture plantations are relatively more vulnerable to environmental disturbance and attacks by pests and pathogens compared with mixed plantations (Huuskonen *et al*., 2021), and they may fail to provide niches to native organisms including plants, animals and microorganisms (Wang *et al*., 2019). The biodiversity of all these groups diversifies ecosystem services and secures habitat persistence via increased tolerance to environmental stress and resilience (Yang *et al*., 2018).

Individual plant species contribute to the relative diversity effect on ecosystem processes through their characteristic functional traits (Diaz & Cabido 2001). Since closely related species have more similar traits, their relative effect can be predicted by phylogenetic relationships among species. Such phylogenetic conservatism phenomenon applies to nearly all plant aboveground and belowground traits including interactions with mutualistic and antagonistic animals and microorganisms (Anacker *et al*., 2014; Gougherty & Davies 2021). Several studies have indicated that plant communities with diverse functional traits promote soil biodiversity and ecosystem functioning relatively more than species-rich communities with low functional diversity (Cadotte *et al*., 2011; Ampoorter *et al*., 2020). Associations with root symbiotic mycorrhizal fungi and nitrogen-fixing bacteria are among the most important plant traits from the perspectives of nutrition and ecosystem processes (Martin *et al*., 2017). Plant mycorrhiza types, in particular, determine soil microbial composition and functionality (Bahram *et al*., 2019) as well as soil carbon and nutrient cycling (Read, 1991; Phillips *et al*., 2013; Tedersoo & Bahram 2019).

Plant diversity effects can be most efficiently studied using field experiments that enable avoiding the confounding natural niche differentiation by position in landscape, microsites, soil properties and co-occurring plants (Grossman *et al*., 2018). Botanical gardens, arboreta and well-planned experimental plantations can also be treated as sentinel sites for rapid monitoring of the occurrence and spread of pests and pathogens. Such plantations offer additional information about the population dynamics of antagonist species, their specificity to host plants and relative host plant density dependence (Morales-Rodriguez *et al*., 2019). In principle, such analyses can also be applied to studies of mutualists such as pollinators, mycorrhizal fungi and root-nodulating, nitrogen-fixing bacteria (e.g. Li *et al*., 2021).

Here we report the design and perspectives of a taxonomically inclusive (105 woody species and 11 perennial herbaceous species) MycoPhylo field experiment. This experiment aims to fulfil the following objectives: i) assess the role of tree and shrub taxonomic, functional and phylogenetic diversity and mycorrhiza type diversity on biodiversity and ecosystem processes; ii) determine the effect of plant mycorrhiza type on soil processes and ecosystem function; and iii) offer a sentinel system for monitoring pest and pathogen colonisation and determination of host specificity of antagonist and mutualist communities.

## Materials and methods

The MycoPhylo experiment was founded on former cropland in the Rõhu experimental centre (58.36 °N, 26.52 °E) in April, 2018. Two years previously, the experimental area (1.5 ha) was tilled to a depth of approximately 30 cm and it was densely covered by various grasses and forbs. The site is flat, with max. altitudinal difference of 0.5 m and no trenches. The soils are clay loam, with ca. 30 cm AO horizon, 5 cm of A horizon and a hard, clayey C horizon below 35 cm depth. On average, the site has a soil pH_KCl_ of 6.5, C_total_ of 43.5 g kg^-1^, N_total_ of 3.6 g kg^-1^, phosphate of 0.25 g kg^-1^, K_total_ of 0.39 g kg^-1^, Ca_total_ of 1.50 g kg^-1^and Mg_total_ of 0.20 g kg^-1^ (Tedersoo *et al*., 2021). The mean annual temperature and mean annual precipitation are 5.9 °C and 631 mm, respectively.

The experimental area was divided into a buffer zone and double rows of square experimental plots (4 m x 4 m for trees and bushes, 2 m x 2 m for shrubs and herbs) in the west-to-east direction (Figure 1). The experimental site has a triangular shape and it is surrounded by tree hedges from three sides: *Quercus robur* and *Corylus avellana* (planted in the 1970s) in the south, *Tilia cordata* (planted in the 1970s) in the west and *Tilia platyphyllos* (planted in 2005) in the northeast. The rooting zone of tree rows is well beyond the plantation as determined by root sampling of trees and observations of fungal fruiting bodies. However, these rows of trees provide partial shade to the plantation from the south late in the growing season (September) and from the west in the afternoon (after 2 PM to 3 PM in the growing season). Tree leaves are also blown partly into the experimental area. Spatial analysis is needed to account for these potentially important effects.

**Figure 1.**
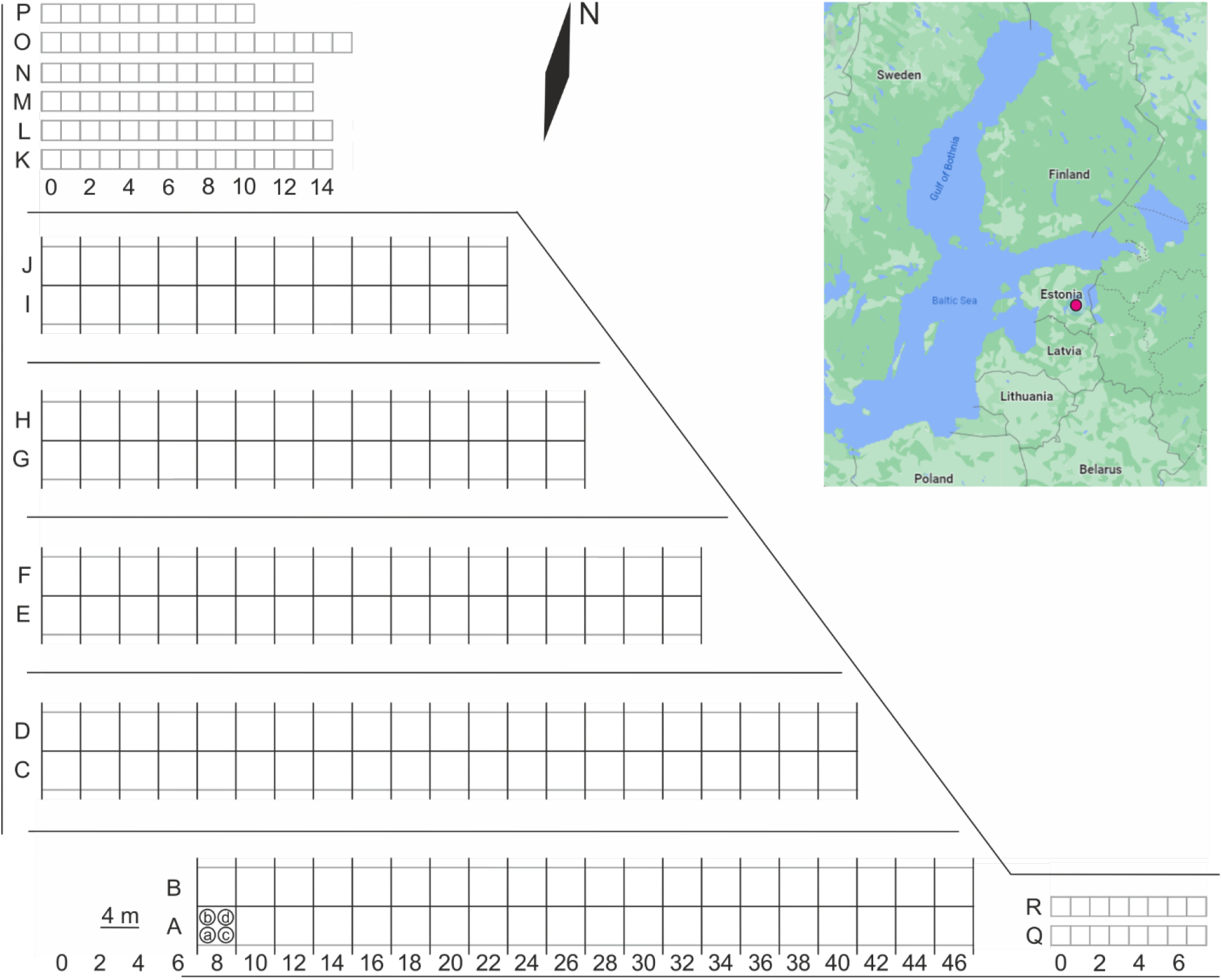
Layout of the MycoPhylo experiment. Cap letters, numbers and encircled small letters indicate row labels, column numbers and positions of individual plants, respectively. Grey lines indicate plot borders; black lines indicate root barriers.

According to the experimental design, the plots harbour a monoculture, diculture (two plants from each two species) or tetraculture (each plant from a different species). For each richness treatment, the four plant individuals belong to one, two or three different mycorrhiza types – arbuscular mycorrhiza (AM), ectomycorrhiza (EcM) or ericoid mycorrhiza (ErM). For tetracultures, the mycorrhiza type with two plant species was randomly selected. Each mycorrhiza type was also represented by all mono-, di- and tetracultures. There were five species from each mycorrhiza type that were used with four replicates in monocultures and mixed treatments (EcM: *Betula pendula* Roth, *Populus tremula* L., *Quercus robur* L., *Salix caprea* L., *Tilia cordata* Mill.; AM: *Acer platanoides* L., *Fraxinus excelsior* L., *Ulmus laevis* Pall., *Prunus padus* L., *Sorbus aucuparia* L.; ErM: *Empetrum nigrum* L., *Vaccinium vitis-idaea* L., *V. macrocarpon* Aiton, *Calluna vulgaris* (L.) Hull and *Rhododendron* sp.). Plant combinations in dicultures (n=30) and tetracultures (n=40) were assigned randomly without replacement but identical combinations were avoided. Four monoculture replicates were also included for *Picea abies* (L.) H.Karst, *Pinus sylvestris* L., *Pseudotsuga menziesii* (Mirb.) Franco (EcM conifers), *Thuja occidentalis* L., *Juniperus communis* L. *and Taxus baccata* L. (AM conifers). Multiple other native and non-native trees, bushes, shrubs and herbal species of various mycorrhiza types were grown in unreplicated monocultures to add power to analyses of the phylogeny effect. The shrubs and bushes were planted in 2 × 2 m monoculture plots at 1 × 1 m intervals. A total of 12 perennial herb and shrub individuals from 15 species (including the putatively non-mycorrhizal species *Carex muskingumensis, Armoracia rusticana, Luzula Pilosa and Lupinus polyphyllus;* Soudzilovskaia *et al*., 2022) were planted as unreplicated monocultures in 2 × 2 m plots (three rows separated by 20 cm with a 55 cm distance between them). Initially, 265 plots comprising 132 plant species were established. As of May 2022, 237 plots and 116 plant species survived (i.e., at least three individuals were alive).

The planting stock originates from various nurseries in Estonia and Latvia and one nursery in Poland. We also replanted saplings from Estonian woodlands and forests if these species were not grown in nurseries or were too small. Accordingly, seedlings and saplings were excavated and transported to the plantation site with roots embedded in soil (5-50 l depending on plant size; originating from forests and some nurseries), potted (0.5-10 kg soil; nurseries only) or bare-rooted (nurseries only; Table S1). Planting was performed manually in April and May 2018. Dead plants were replaced in August 2018 and May 2019.

Three monocultures were planted additionally in May, 2019 (Table S1). Plants that died in summer 2019 were occasionally replaced by other plants in the experimental plantation to secure four focal plants in the plots and eliminate plots with poor plant survival (11 plots). All plants received 5-10 l of water at planting. In 2018, all plants were watered weekly with 5-10 l of water because of a severe drought from early June to August. A larger area around the experiment was fenced to prevent damage by hares and larger herbivores.

The plots are mowed monthly with a small tractor from May to September, and manually biweekly around tree trunks. Shrubs and herbs are weeded manually without disturbing the soil to remove competing herbs and avoid damage to focal plants. In April 2022, the 4 × 4 m tree plots were separated from each other using a polycarbonate (4 mm diam.) barrier placed at 0 to 40 cm depth to prevent root ingrowth from neighbouring plots (Fig. 1). All sides of the experimental area were similarly separated to minimise potential belowground effects from the neighbouring tree rows.

At the time of planting, a mixed sample of soil and fine roots (roughly 1:1 vol.) was collected from four individuals of each plant species. These four samples were pooled, dried in a drying cabinet at 35 °C and subjected to DNA extraction from 0.25 g of bead-homogenised material using the Soil UltraClean DNA Isolation kit (MoBio, Solana Beach, CA, USA) following the manufacturer’s instructions. The DNA samples were maintained at -20 °C until use. We analysed all eukaryotes from these samples using the tagged universal primers ITS9mun and ITS4ngsUni as described in Tedersoo *et al*. (2021). The samples will reveal the initial pre-planting microbiome, enabling us to track the mutualistic and antagonistic organisms. The control rhizosphere samples were obtained from six local plant species. Five additional control samples of soil and roots were collected from the planting area and surrounding area in 2019 following the Global Soil Mycobiome consortium design (data released in Tedersoo *et al*., 2021) for analysis of soil microbiome and chemical properties. In April 2022, we performed an initial screening of annual (for 2021) and total height growth indicative of productivity. At the same time, we systematically evaluated the health status of trees including damage by frost, pathogens, pests and rodents. These measurements will be performed in April each year.

The phylogenetic relationships between cultivated plants were obtained from the phylogenetic tree of land plants generated by Zanne *et al*. (2014) using the ‘keep.tip’ function in the ape package. The phylogenetic diversity of each plot was calculated with the ‘pd.calc’ function in the caper package of R, using the total branch length (TBL) method (Orme *et al*., 2013). The phylogram and Newick-formatted tree are given in Figure S1 and Item S1, respectively.

## Results and Discussion

The MycoPhylo experiment is specifically designed to test plant phylogeny effects on plant and soil microbiome and ecosystem function from the mycorrhiza perspective. The main benefit of this experiment is the wide variety of taxa (105 woody plant species from 36 families), the array of life forms (trees, bushes, shrubs and herbs) and the large number of experimental units (237 plots). The taxonomic breadth is important, because in both regular and phylogeny-aware analyses, species and higher taxonomic groups contribute to the number of degrees of freedom, respectively (Orme *et al*., 2013). Plant phylogeny was an important predictor in the study of host effects on foliar fungal endophyte communities in the BiodiversiTREE experiment (Griffin *et al*., 2019). In comparison, another taxonomically inclusive BEF-China experiment comprises 40 tree species and 20 shrub species from 33 families (Bruelheide *et al*., 2014). In the BEF-China experiment, all 40 tree species are represented by replicated monocultures, whereas 21 woody plant species were initially represented by replicated monocultures in the MycoPhylo experiment (*Calluna vulgaris* monocultures perished).

Our experiment was also designed to assess the effect of mycorrhiza types of plants and mycorrhiza type richness of plant community. The belowground plant traits mycorrhiza type and N-fixing associations play a significant role in ecosystem functioning (Phillips *et al*., 2013; Tedersoo & Bahram, 2019) and soil microbiome composition (Bahram *et al*., 2020). MycoPhylo is the first plant diversity experiment to include ErM plants in sufficient replicates (4 surviving species with replicated monocultures and six species with a single monoculture replicate). Notably, all ErM plants are shrubs or small bushes; to account for this life form bias, we also included multiple shrubby EcM and AM plant species. We also opted for a high degree of phylogenetic diversity of EcM plants – initially 32 species from 18 genera and 7 lineages (sensu Tedersoo & Brundrett 2017; 29 species from 16 genera and 5 lineages survived) – because members of different EcM plant lineages differ strongly in their ecophysiological traits (Koele *et al*., 2012). Design of certain other experiments (e.g. BiodiversiTREE; BEF-China; Macomer, van de Peer *et al*., 2018; and MyDiv, Ferlian *et al*., 2018) allows testing mycorrhiza type effects, but these experiments use EcM trees belonging to a maximum three EcM plant lineages. In the MyDiv experiment, all tree species have been reported as dual mycorrhizal, where EcM colonisation of the predicted AM plants often exceeds that of predicted EcM plants (Heklau *et al*., 2021), challenging the mycorrhiza type comparisons (if true). Furthermore, these experiments do not use root exclusion mechanisms, which may strongly blur measurements of soil biodiversity and function. For example, Singavarapu *et al*., (2022) reported that on average, 12-17% of fungal reads in AM-AM plant plant neighbours represented EcM fungi from other surrounding trees. Preliminary results from other tree diversity experiments indicate that EcM trees have a relatively lower growth rate compared with AM trees and their mixture has no synergistic effects on productivity (Ferlian *et al*., 2018). Mixing with other trees reduces insect damage to AM trees but not EcM trees (Ferlian *et al*., 2021). EcM Fagales harbour a relatively lower diversity of foliar fungal endophytes compared with AM plants (Griffin *et al*., 2019). The diversity of neighbouring trees reduced the proportion of specialists and promoted foliar fungal diversity in young saplings (Weissbecker *et al*., 2019) but not in older saplings (Li *et al*., 2021). AM trees support more diverse soil fungal communities than EcM trees, but there is no difference in bacterial richness and functionality (Singavarapu *et al*., 2022).

The occurrence of monocultures of both native and introduced trees and shrubs allows us to estimate the relative effect of non-native plants on soil microbiome and functionality. We refrained from using introduced plant species (except *Vaccinium macrocarpon* and *Rhododendron* sp.) in mixed plant treatments to avoid confounding the effects of diversity and the presence of non-native species. In this respect, our experiment resembles arboreta and botanical gardens, where species from multiple origins are planted in an aggregated manner. Such design benefits the experiment for inclusion in a system of sentinel plantations and recording local pests and pathogens in introduced plants and vice versa.

The MycoPhylo experiment is one of the northernmost plant biodiversity experiments. To the best of our knowledge, only a forestry field experiment in Satakunta (5 species) in Finland occurs in the more northern, boreal forest biome (Vehviläinen & Koricheva 2006). Most other experiments are located in the warm temperate zone in European countries and USA (Verheyen *et al*., 2016). Therefore, these northernmost experiments offer valuable information about the BEF effects in the boreal and hemiboreal climate. From the mycorrhiza type perspective, experiments are also needed in tropical regions, because limiting nutrients and plant adaptations likely differ.

While the MycoPhylo experiment exhibits unique features, it also has several limitations. First, the small size of plots (4 m x 4 m and 2 m x 2 m) does not allow the development of a forest microclimate and is irrelevant from the perspectives of forestry as well as bird and mammal studies. The entire plots are subjected to strong edge and neighbour effects, with a negligible exterior to interior gradient. These effects are partly ameliorated by our focus on soil habitats and the establishment of plastic root barriers. Although these barriers minimise ingrowth of root and fungi from neighbouring plots, they may alter the drainage and limit migration of free-living soil organisms. Second, the four plant individuals per plot limit the biodiversity gradient to 1-4 species and make plots vulnerable to the loss of any single plant individual. Although we substituted dead plants in early years, higher mortality may be expected in the future due to exceptionally cold winters, dry summers or pathogen outbreaks (e.g. the ash dieback agent *Hymenoscyphus fraxineus* in *F. excelsior* that caused high mortality in 2018; Table S2). Up to 18 co-occurring woody plant species have been planted per plot in tree diversity experiments, but most use four species for the highest-diversity treatment (Grossman *et al*., 2018).

In conclusion, the MycoPhylo experiment is particularly useful for addressing questions from the mycorrhizal type and plant phylogeny perspectives. This experiment has joined the TreeDivNet network and welcomes collaborative research in testing hypotheses in biodiversity-ecosystem function and species effects in large-scale metastudies.

## Supporting information

Item S1

## Acknowledgements

We are indebted to multiple students and colleagues who helped with transporting and planting seedlings and saplings. The project was financed by the EEA Financial Mechanism Baltic Research Programme (EMP442), Novo Nordisk Fonden (NNF20OC0059948), EU Regional Development Fund (AnaEE Estonia 4.01.20-0285) and Estonian Research Council (PRG1615).

## Author contributions

LT, KL and RD designed the experiment, all authors maintained the experiment and performed measurements; LT wrote the paper with input from all coauthors.

## Data availability

Information about plot locations and plants is available in the online supplement. FastQ files of rhizosphere eukaryotes are available from the Short Read Archive under accession XXXXXX.

## Supplementary materials

**Figure S1.** Phylogram of plant species used in the MycoPhylo experiment.

**Item S1.** The Newick-formatted tree of plant species used in the MycoPhylo experiment.

**Table S1.** Information about plant species used in the MycoPhylo experiment including planting and survival details and codes used in molecular analyses.

**Table S2.** Information about plots used in the MycoPhylo experiment.

