## Supplementary figures and images for "MycoPhylo experiment reveals how mycorrhiza types and phylogenetic relationships affect soil biodiversity and functioning"

### Figure S1

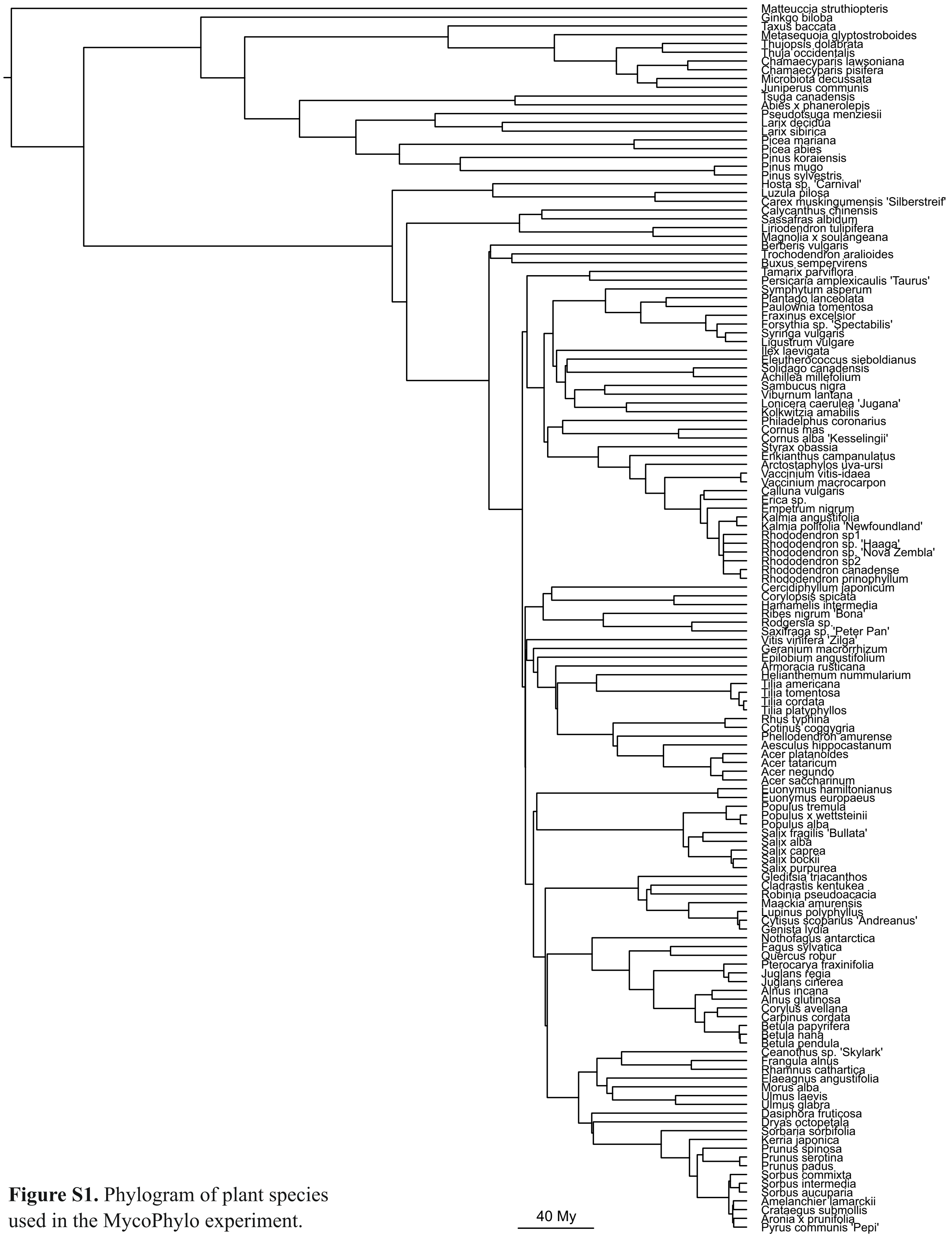


**Figure S1.** Phylogram of plant species used in the MycoPhylo experiment.
